# A Genetically Encodable and Chemically Disruptable System for Synthetic Post-Translational Modification Dependent Signaling

**DOI:** 10.1101/2022.05.29.493928

**Authors:** Jeffrey B. McMahan, John T. Ngo

## Abstract

We describe an engineered “writer/reader” framework for programming post-translational control into synthetic mammalian signaling proteins. In this approach, a bacterially-derived biotin protein ligase (BirA) was used as a “writer” element for the modification of artificial receptors and transcription factors containing a biotin acceptor peptide (AP) fusion tag. To enable modification events to transmit biochemical information, we designed encodable “reader” modules using sequences from a biotinamide-binding antibody. Proteins fused to reader domains were able to interact with AP-tagged polypeptides in a biotinylation-dependent manner, and control over the timing and extent of these interactions could be modulated through both genetic and chemically-based strategies. Genetic and cell-specific control over AP-reader module interactions was achieved via regulated BirA expression, and the interaction states of both intra-and inter-cellular complexes could be modulated with biotinamide-based and bioorthogonally-functionalized compounds. The utility of this approach was demonstrated by installing post-translational and chemogenetic control into synthetic Notch (“SynNotch”)-based systems.

## INTRODUCTION

A central goal in synthetic biology is to program cellular systems to carry out sophisticated tasks resembling those that are found in nature. Toward this goal, researchers have devised strategies in order to regulate biological processes, with the goal of adapting cells and their assemblies for customized applications in science, medicine, and biotechnology. Over the past two decades, these efforts have led to powerful strategies for controlling gene expression, signal transduction, and cell-cell communication^1,2^. In many cases, these systems have involved the construction of synthetic multidomain proteins, in which customized polypeptide modules are fused together in order to mimic the functions and organization of endogenous signaling machinery^3^. A prominent example of success in this approach has been the design of artificial signaling systems based on the Notch receptor, referred to as synthetic Notch (“SynNotch”) proteins^4–6^. Like natural Notch^7,8^, SynNotch receptors are transmembrane signaling proteins composed of modular units, including an extracellular ligand-recognition element and an intracellular transcriptional regulatory domain. Through a mechanism that is referred to as “mechanical allostery,”^4^ the binding of ligands to the receptor extracellular domain (ECDs), in combination with the delivery of sufficient mechanical tension^4,9–11^, results in the proteolytic release of the intracellular domain (ICD) such that it is able to translocate to the nucleus for the regulation of gene expression. By substituting ECD and ICD components surrounding a conserved SynNotch core segment – a sequence composed of the natural receptor’s negative regulatory region (NRR) and transmembrane domain (TMD) – one is able to generate customized receptors, with which cells can be endowed with user-specified sense-and-respond capabilities.

In recent years, the SynNotch strategy has been implemented in various applications, ranging from basic investigations of Notch signaling and NRR function^4,12^, to the programming of synthetic cells for immunotherapy and tissue engineering applications^5,6,13^. Strategies to regulate synthetic receptors have also been developed, including those mimicking natural Notch regulatory mechanisms, such as *cis-*inhibition and synthetic lateral inhibition programs^13–15^. However, despite the versatility of the current framework, SynNotch designs lack the sophisticated control features that are imparted to endogenous Notch components via post-translational protein modifications.

In natural systems, post-translational modifications (PTMs) serve as critical modulators of protein functions in countless biological processes. For example, the modification of proteins via covalent chemical additions, or by cleavage of polypeptide backbones, can result in rapid alterations in protein properties, including changes in their three-dimensional structures, interaction states, half-lives, and subcellular localizations^16,17^. In the context of Notch signaling, the modification of receptor components provides a crucial layer of regulation through which the diverse pleiotropic and context-dependent outcomes of signal transduction are manifested *in vivo*^18–21^. Emblematic of such mechanisms is the glycosylation of the Notch ECD, which occurs within the endoplasmic reticulum (ER) and Golgi apparatus in order to control the binding-specificities of receptors and ligands at the cell surface^19,22,23^. These modifications contribute to the spatiotemporal regulation of Notch-mediated cell-cell communication during multiple developmental processes, including the control of tissue boundary formation, as well as that of vertebrate segmentation. In addition to extracellular PTMs, the modification of intracellular components is also critical to Notch signaling control, including the phosphorylation of the native ICD, which serves to restrict the duration and magnitude of Notch signaling responses by tuning the half-lives of liberated ICDs within the nuclear compartment^24,25^. Mutations resulting in the alteration of Notch PTMs have been linked with multiple disease states, including various human cancers, thus underscoring the critical nature of these modifications in the regulation of the endogenous pathway. Seeking to impart similar control into engineered signaling programs, we thus set out to design a synthetic strategy for encoding PTM-mediated regulation into SynNotch and its signaling components.

## RESULTS

### A Biotinylation-Sensitive Synthetic Notch

In designing a synthetic PTM-control strategy, we outlined criteria that would be required to achieve versatile and orthogonal control over SynNotch proteins. In an ideal scenario, one would be able install a PTM site into any desired signaling component via direct genetic fusion with a minimal substrate tag. In addition, in order to facilitate the PTM of the protein, a modifying enzyme capable of catalyzing selective and covalent modification of the tag should also be in hand. Both the tag and enzyme should be absent from the endogenous proteome of the engineering chassis, as to limit crosstalk between synthetic and endogenous signaling components. Furthermore, the enzyme should modify only its designated (tagged) synthetic protein targets within the context of the engineered cell. To achieve predictable and tunable levels of protein modification, the enzyme should have a strict and well-defined substrate specificity, and the ability to function within both extra-and intra-cellular compartments would be an additional desirable feature. Importantly, in order to have utility in cell engineering applications, the synthetic PTM event must be able to transmit synthetic biochemical information in some way.

In considering systems that could satisfy these design criteria, we turned our attention to biotin protein ligase (BirA) from *Escherichia coli*, which catalyzes the covalent chemical attachment of biotin to lysine side chains encoded within its substrates. Although BirA’s natural substrate (biotin-carboxylase cargo protein, BCCP)^26^ is composed of 156 amino acids, short polypeptide substrates for the enzyme have been identified, including a 13 amino acid sequence referred to as the biotin acceptor peptide (AP)^27^, as well as a related 15 amino acid sequence known as “AviTag”^28^. Given the orthogonal biotinylation activity of BirA within multiple mammalian cell compartments^29–32^, we reasoned that the bacterial ligase could be implemented as an orthogonal “writer” protein for the selective modification of synthetic signaling components containing AP or AviTag sequence fusions. To test this possibility, we proceeded by designing and testing a biotinylation-sensitive SynNotch receptor.

### Characterization and validation of Acceptor Peptide (AP)-SynNotch

A biotinylation-sensitive synthetic Notch (SynNotch) was constructed by fusing the biotin acceptor peptide (AP) to the extracellular region of an anti-GFP containing receptor, generating “AP-SynNotch.” In this design, we anticipated that the ligand-specificity of the receptor could be regulated in a post-translational manner, enabling its recognition of biotin-binding proteins following modification by BirA (**Figure 1a**). In initial experiments, we sought to confirm that AP-SynNotch could be selectively modified in cells containing luminal BirA constructs, and that the receptor could be correctly processed and trafficked to the cell surface following biotinylation. Immunoblot analyses of cells expressing either an endoplasmic reticulum (ER)-, or Golgi-targeted BirA (BirA-KDEL and GalT-BirA, respectively), showed highly-specific modification, with receptor components appearing as the only streptavidin-reactive bands beyond that of endogenously biotinylated proteins (**Figure 1b, Supplementary Figure 1a-b**). Notably, signals corresponding to modified versions of both full-length (77 kDa) and S1-cleaved (43 kDa) receptor fragments were observed, with detection of an extracellularly-encoded myc epitope providing confirmation of the identity of these bands. Thus, post-translationally modified AP-SynNotch can be correctly processed via furin cleavage within the Golgi to produce a mature, heterodimeric receptor^33^. Because more efficient biotinylation was observed using BirA-KDEL, as compared to GalT-BirA, we proceeded with the ER-localized enzyme in subsequent analyses.

**Figure 1.**
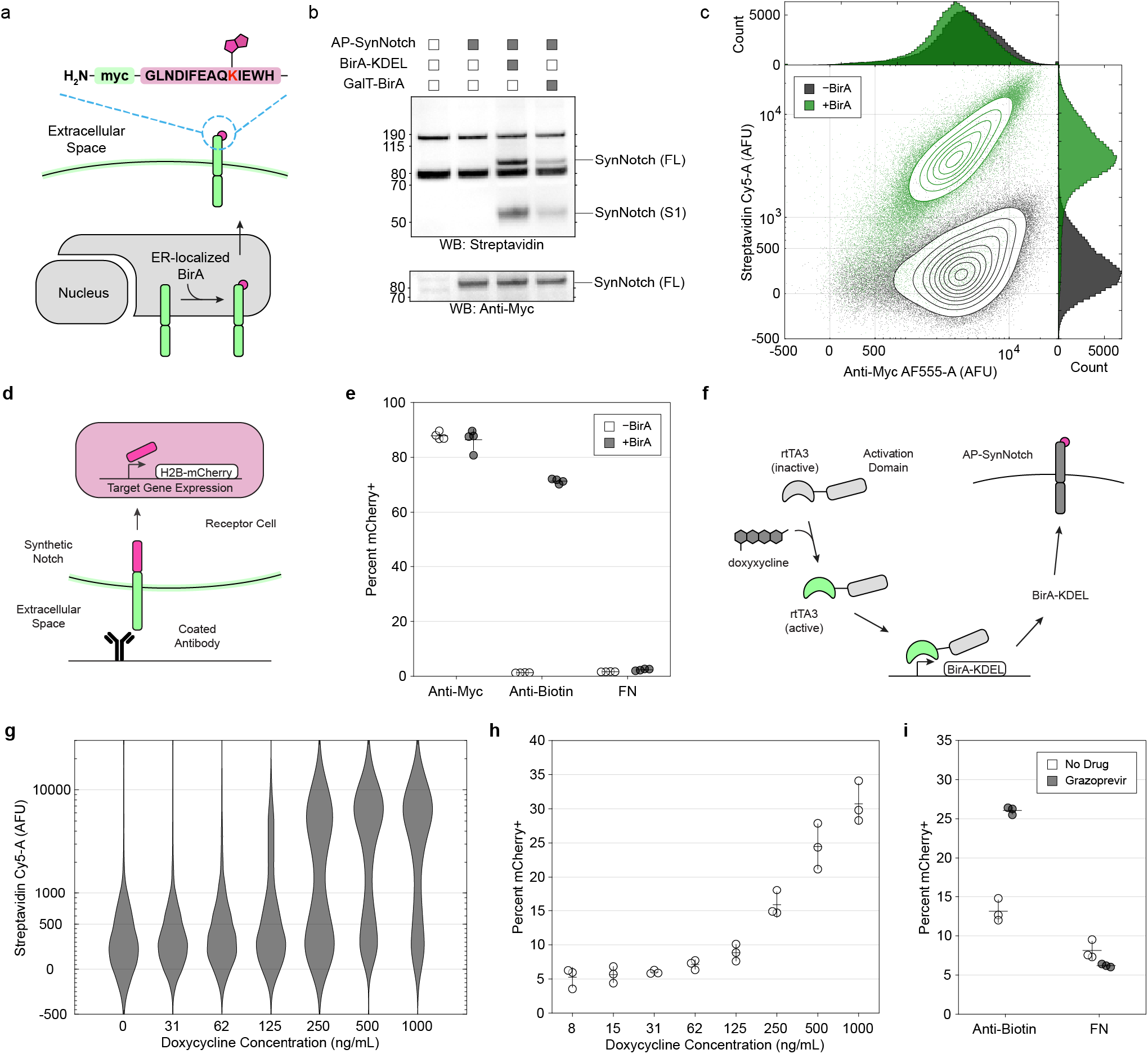
Biotinylation and signaling activity of AP-SynNotch. **a**. Schematic depicting the post-translational biotinylation of AP-SynNotch by BirA-KDEL. **b**. Western blot analysis of AP-SynNotch in transfected HEK293 cells containing coexpressed BirA-KDEL, GalT-BirA, or without BirA. Fragments corresponding to full-length (FL) AP-SynNotch and its furin/(S1)-cleaved extracellular fragment are indicated. Lysate from non-transfected HEK293 cells was used to identify endogenously biotinylated protein bands. **c**. Flow cytometry analysis of cell-surface localized AP-SynNotch in BirA expressing (+BirA) and non-expressing cells (-BirA). Non-permeabilized cells expressing AP-SynNotch were stained using anti-myc antibody and streptavidin in order to determine the surface presentation levels of the unmodified and modified receptors. The BirA positive well included 146,160 cells, and the BirA negative well included 138,617 cells. **d**. Schematic depiction of the activation of AP-SynNotch using an immobilized anti-biotin IgG-based ligand. The nuclear translocation of the ICD and its induction of reporter (H2B-mCherry) expression is shown. **e**. Quantification of signaling-induced reporter expression in AP-SynNotch cells stimulated with either anti-biotin IgG, or anti-myc IgG as a positive control. Reporter levels are shown from cells with and without coexpressed BirA-KDEL. Signals from ligand-stimulated cells are shown in comparison to control wells coated with fibronectin (FN) only. Error bars represent one standard deviation, n = 4 wells. **f**. Schematic depicting the transcriptional control of BirA-KDEL through doxycycline induction of rtTA3. **g**. Flow cytometry measurement of biotinylated AP-SynNotch on the surface of cells encoding a doxycycline-inducible BirA-KDEL construct (TRE3G-BirA-KDEL) and expressing rtTA3. Cells treated with varying concentration of doxycycline were analyzed by cell surface staining using streptavidin-Cy5. Wells included approximately 10,000 cells per condition. **h**. Quantification of signaling-induced reporter expression in TRE3G-BirA-KDEL cells containing rtTA3 and stimulated with fixed amounts of immobilized anti-biotin IgG and varying concentrations of doxycycline. Error bars represent one standard deviation, n = 3 wells. **f**. Reporter expression levels from AP-SynNotch/TRE3G-BirA-KDEL cells in which 5 μM grazoprevir was used to induce BirA-KDEL expression. Error bars represent one standard deviation, n = 3 wells.

Given that cell surface localization is required for the detection of extracellular ligands, we next asked whether biotinylated AP-SynNotch could be trafficked to the plasma membrane. Staining of non-permeabilized cells using dye-conjugated streptavidin confirmed the presentation of biotinylated receptors on the surface of transfected HEK293 cells (**Supplementary Figure 1c**). In addition, labeling with anti-myc antibody showed comparable levels of surface reactivity between cells that expressed BirA-KDEL and non-expressing controls (**Figure 1c, Supplementary Figure 1d**). Thus, biotinylation did not appear to deter the trafficking efficiency of AP-SynNotch, as compared to its unmodified counterpart. These results, together with those described above, confirm that AP-SynNotch can be selectively and efficiently biotinylated during secretory transport, and that the modified receptor is correctly processed and trafficked by endogenous cellular machinery prior to its presentation at the plasma membrane.

### Conditional signaling in response to a PTM-specific ligand

We next evaluated the signaling properties of AP-SynNotch, asking whether biotinylation of the receptor altered its activity in response to different synthetic ligands. Here, a receptor containing a Gal4-VP64 intracellular domain (ICD) was evaluated using HEK293 cells containing a stably-integrated reporter gene (UAS:H2B-mCherry). Following transfection with receptor-encoding DNAs, cells were grown on culture surfaces containing immobilized ligands in order to stimulate the activation of AP-SynNotch (and thus nuclear translocation of the Gal4-VP64 ICD). After overnight growth in the presence of each ligand, we quantified signaling responses using flow cytometry, comparing the extent of reporter expression between BirA-KDEL expressing and non-expressing cells (**Figure 1d**).

In initial analyses, we evaluated the background activity of AP-SynNotch, asking whether its biotinylation by BirA-KDEL altered signaling quiescence in the absence of corresponding ligands. Quantification of reporter levels from ligand-untreated cells showed similar degrees of background activity between BirA-KDEL containing and non-expressing controls (**Figure 1e**). Thus, the coexpression of AP-SynNotch and BirA-KDEL did not appear to induce discernible levels of ligand-independent activity. To confirm the inducibility of the receptor, we also analyzed cells that were stimulated with immobilized anti-myc IgG, a positive control ligand (**Supplementary Figure 1e**). Measurement of these cells revealed strong levels of signaling-induced reporter expression, with H2B-mCherry intensities that were closely-matched between BirA-KDEL expressing and non-expressing cells. Thus, the expression of BirA-KDEL did not appear to prevent, nor limit the inducibility of AP-SynNotch in response to a PTM-independent ligand.

In contrast to our results using anti-myc IgG, stimulation of cells with a biotin-specific ligand (anti-biotin IgG) resulted in divergent signaling responses (**Figure 1e, Supplementary Figure 1f**). Whereas cells that expressed BirA-KDEL exhibited strong signaling activity (with reporter levels nearing those that were induced using anti-myc IgG), cells that lacked BirA-KDEL expression exhibited only background amounts of H2B-mCherry, similar to that of ligand-untreated controls. Together, these data confirm the ligand-dependence of AP-SynNotch in both its unmodified and biotinylated states. Furthermore, these results provide evidence that BirA-KDEL is able to confer new ligand-recognition capabilities to AP-SynNotch, enabling its detection of biotin-binding ligands in a PTM-dependent manner.

### Control over receptor PTM and signaling activity

In natural systems, the PTM of Notch receptors is tightly regulated in order to achieve precise control over developmental processes such as cell patterning, boundary formation, and tissue morphogenesis^22,34,35^. In certain systems, this regulation is achieved via dynamic and spatiotemporally-restricted expression of Notch-modifying glycosyltransferases genes^36,37^. Seeking to gain similar synthetic control, we next asked whether regulated BirA-KDEL expression could be used to fine-tune the extent of biotinylation, and as a result, the degree of signaling responses to biotin-binding ligands.

In an initial analysis, we confirmed that graded levels of receptor biotinylation could be facilitated by transfecting cells with varying amounts of BirA-KDEL plasmid (**Supplemental Figure 1g**). Desiring to gain greater control, we next asked whether more precise fine-tuning could be achieved by regulating the amount of ligase expression at the level of gene transcription. To test this possibility, we generated a stable cell line in which BirA-KDEL expression was placed under control of the TRE3G promoter. In cells where the tetracycline transactivator-3 (rtTA-3) was used to control TRE3G transcription, both the extent of receptor modification, as well as cell signaling responses to anti-biotin IgG, could be tuned in a dose-dependent manner using doxycycline (**Figure 1f-h**).

In addition to rtTA-3, similar control was achieved using an inducible transcription factor based on the LInC (for “Ligand Inducible Connection”) strategy^38,39^. Here, a transcription factor that is responsive to viral protease inhibitors – specifically those targeting the Hepatitis C virus (HCV) NS3 *cis*-protease – was exploited in order to achieve drug-mediated control via grazoprevir, a clinically approved antiviral compound. In this approach, BirA-KDEL expression was regulated using a fusion protein in which the *cis*-protease was inserted between TetR and VP64 domains (TetR-NS3-VP64). Accordingly, treatment with grazoprevir resulted in elevated AP-SynNotch signaling in response to immobilized anti-biotin IgG (**Figure 1i**). Together with our results using doxycycline and rtTA3, these data confirm that AP-SynNotch biotinylation and its signaling responses can be controlled via BirA-KDEL modulation. Furthermore, the results provide evidence that not only the signaling specificity of AP-SynNotch, but also its signaling magnitudes, can be defined in a drug-inducible and PTM-dependent manner.

### An encodable biotinamide-binding ligand

Although IgG-coated surfaces provide a convenient way to quickly validate receptor designs, our goal in this work was to construct synthetic signaling systems that closely resemble those found in nature. Thus, we next sought an encodable biotin-binding ligand, anticipating that such a protein could be utilized in order to generate a synthetic and PTM-dependent intercellular signaling pathway. Hypothesizing that such a ligand could be made by expressing a biotin-recognizing domain as a cell surface protein, we examined known biotin-binding proteins to identify a suitable recognition element. Because natural Notch ligands and receptors interact in a one-to-one stoichiometry^14^, we were particularly interested in sequences that could be readily converted into monomeric/monovalent binding domains. Thus, although streptavidin-and avidin-derived sequences were considered, their multimeric natures motivated us to explore an alternative scaffold for the design of a reader domain.

Given our success in using anti-biotin IgG as a model ligand, we asked whether a biotin-binding immunoglobulin could be used to create an encodable PTM-recognition module. Conveniently, previous work has described the structure and sequence of a monoclonal, humanized antibody possessing low-nanomolar binding affinity against biotinamide^40^. Biotinamide is derivative of biotin that is formed following its attachment to proteins, and it also is generated upon modification of AP by BirA – thus, we tested whether a single-chain variable fragment (scFv) based anti-biotinimide could be used to generate a functional monomeric reader element. Using sequence information available within the Protein Data Bank (PDB) we designed an anti-biotinamide scFv and generated a candidate ligand by fusing the fragment to a transmembrane domain (TMD). Because endocytosis is a known requirement in the activation of natural Notch^12^, the intracellular segment of Delta Like (DLL)-1 (a native Notch ligand) was included as the ligand’s cytoplasmic domain, and a SNAP-tag was inserted within its extracellular region of the protein to permit its cell surface labeling and visualization. The resulting sequence, dubbed “anti-bio-ligand,” was then tested for its expression and *trans*-cellular signaling activity (**Figure 2a**)

**Figure 2.**
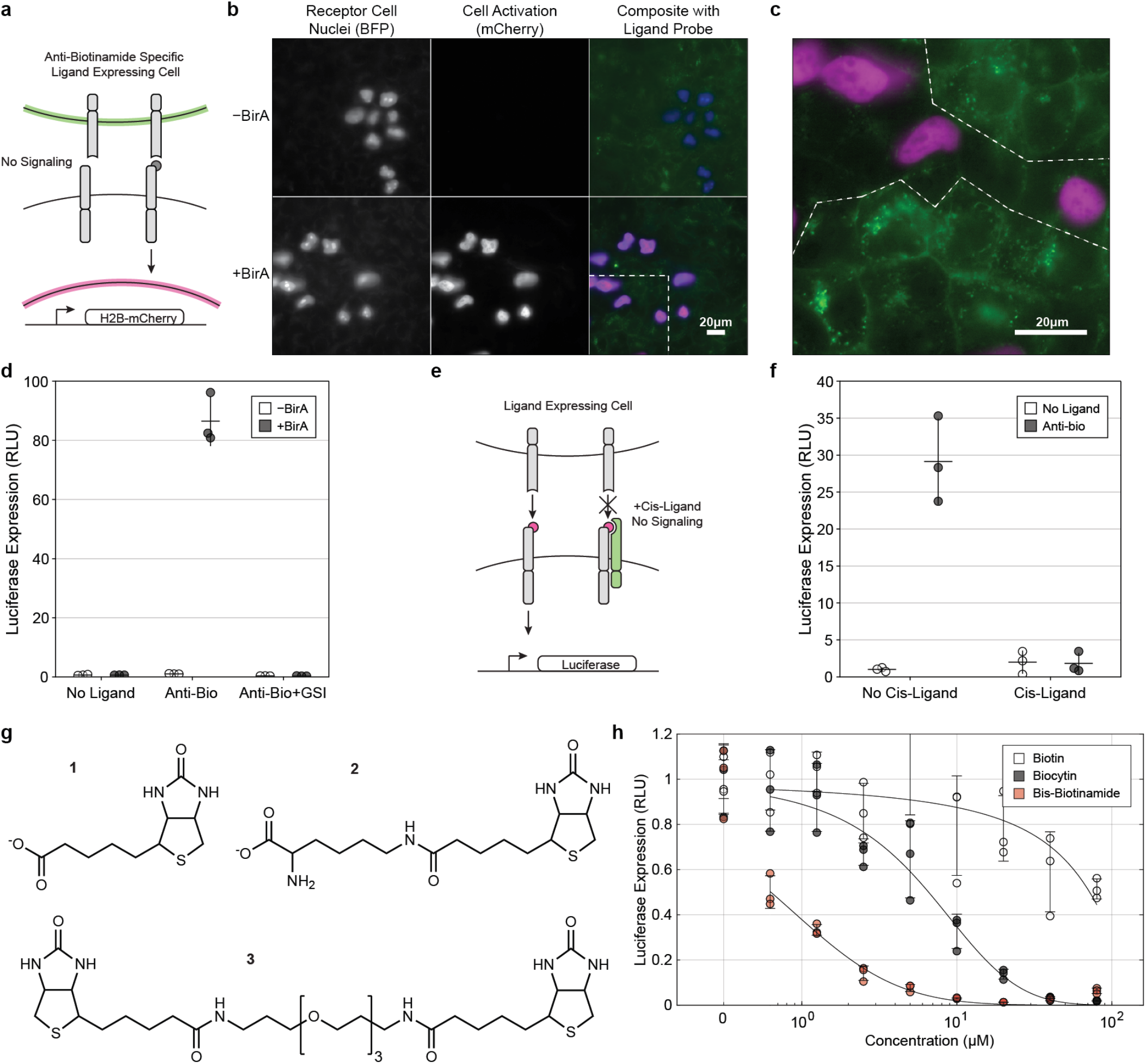
Cell-mediated trans-activation and chemical regulation of signaling from biotinylated AP-SynNotch receptors. **a**. Schematic depicting the biotinylation-dependent *trans*-cellular activation of AP-SynNotch by anti-bio-ligand expressing sender cells. **b**. Representative fluorescence images of cocultures containing either BirA-KDEL expressing or non-expressing receiver cells. AP-SynNotch expressing receiver cells were labeled using a nuclear-BFP marker (blue), and sender cells were identified via direct dye-labeling of anti-bio-ligand using SNAP-Surface-AF647 (green). Expression of the fluorescent reporter protein H2B-mCherry (red) was observed in a contact-and BirA-dependent manner. **c**. Magnified image from inset shown in (b) with overlay indicating cell boundaries. Dashed white line indicates interfacial boundary between sender and receiver cells. **d**. Quantification of *trans*-cellular activation in cocultures using receiver cells bearing a firefly luciferase (fLuc)-based reporter construct (UAS-fLuc). Reporter levels are shown from receiver cells with and without coexpressed BirA-KDEL. Signals are shown in comparison to those from cocultures treated with DAPT, a gamma secretase inhibitor (GSI). Error bars represent one standard deviation, n = 3 wells. **e**. Schematic depicting biotinylation-dependent inhibition of AP-SynNotch in *cis*. **f**. Quantification of UAS-fLuc reporter activation from cocolutres containing *cis*-inhibited receiver cells. Error bars represent one standard deviation, n = 3 wells. **g**. Structures of chemical ligands used competitive inhibitors of transcellular signaling: (1) biotin. (2) biocytin. (3) bis-biotinamide **h**. Dose-response analysis of trans-cellular signaling inhibition using biotin, biocytin, and bis-biotinamide. Error bars represent one standard deviation, n = 3 wells.

In initial analyses, we determined whether anti-bio-ligand could be presented on the cell surface and subsequently internalized into endocytic compartments. Using a “pulse-chase” analysis, we labeled surface-accessible ligand copies during a “pulse” in which cells were treated with cell-impermeant, SNAP-tag reactive dye (SNAP-Surface-AF647). Following a brief labeling period, cells were exchanged into fresh media for a 30 m “chase,” during which the fates of the dye-labeled ligand copies could be tracked. Imaging of dye-labeled cells and cocultures confirmed the surface presentation of anti-bio-ligand, and the detection of dye-labeled ligand copies within internalized punctae verified its ability to be retrieved from the plasma membrane (**Supplementary Figure 2a**). Thus, anti-bio-ligand exhibits the basic cell surface and endocytic properties that are required for *trans*-cellular signal transduction.

Next, to determine whether anti-bio-ligand can facilitate cell signaling in *trans*, we generated a stable line of anti-bio-ligand expressing “sender” cells, which we then used with AP-SynNotch expressing “receiver” cells in a coculture assay. Here, *trans*-cellular signaling activity was evaluated by inspecting receiver cells for the expression of a signaling-induced (Gal4-dependent) reporter gene. Using receiver cells containing a fluorescence-based reporter construct (UAS:H2B-mCherry), we carried out spatial analysis in which cocultures were visualized using fluorescence microscopy. In order to distinguish between individual cell types, we marked receiver cells with a constitutive nuclear-BFP label prior to cell mixing, and we identified sender cells by direct dye-labeling with SNAP-Surface-AF647 *in situ*. In cocultures containing BirA-KDEL expressing receivers, we observed strong H2B-mCherry expression in areas where sender cells and receiver cells were positionally juxtaposed (**Figure 2b-c**). Importantly, reporter expression was not detected within single cultures of BirA-KDEL receivers grown alone (without senders), nor in cocultures in which receiver cells lacked BirA-KDEL (**Figure 2b, Supplementary Figure 2b**). In addition to microscopy analyses, quantitative luminescence-based measurements using a reporter construct based on firefly luciferase (fLuc, UAS:fLuc) provided further validation of anti-bio-ligand as a signaling-competent and biotinamide-specific ligand (**Figure 2d**). Taken together, these data demonstrate that cells can be endowed with signal-sending capability via the expression of anti-bio-ligand. Furthermore, the data indicate that anti-bio-ligand can be used to carry out synthetic and PTM-dependent juxtacrine signaling with AP-SynNotch expressing receiver cells.

### *PTM-dependent* cis*-Inhibition of AP-SynNotch Receptors*

In natural systems, the signal-receiving capability of Notch-bearing cells can be genetically regulated via coincident expression of receptors and ligands (i.e., “*cis*-inhibition”)^14,41^. In certain cases, the strength and specificity of natural *cis*-interactions is modulated via post-translational control^36^. Recognizing the utility of genetic modulation, we next asked whether the signaling capacity of AP-SynNotch could be regulated via binding to anti-bio-ligand in *cis*. To test this possibility, we coexpressed a ligand variant (in which mCitrine served as the ICD, anti-bio-ligand-mCit) in AP-SynNotch receivers and subsequently tested their signal-receiving capacity against anti-bio-ligand sender cells (**Figure 2e**). Quantification of luminescent reporter levels indicated that AP-SynNotch trans-signaling could be inhibited by the co-expression of anti-bio-ligand-mCit. (**Figure 2f**). Thus, these results show that the signaling receptivity that is conferred by BirA-KDEL can be counteracted via the expression anti-bio-ligand within receiver cells. Collectively, our data suggest that BirA-KDEL and anti-bio-ligand can be combined to encode complex signal-sending and -receiving capabilities between cells.

### Inhibition of signaling via soluble biotinamide molecules

Tight-regulation is a requirement of cell-engineered systems in biomedical applications, and such control can be achieved by combining genetic strategies with external drug-control. Thus, in the elaboration of our designs, we next sought to identify exogenous molecules that could be combined with our encodable components to achieve versatile “chemogenetic” regulation. In an initial strategy, we gained inspiration from therapeutic approaches in which soluble Notch “decoy” polypeptides are used to sequester ligands proteins *in vivo*^42^. In this approach, the binding of decoy fragments to Notch ligands prevents their ability to activate receptors in *trans*. Aiming to mimic this strategy, we tested whether soluble biotinamide-containing molecules could be implemented in a similar manner, anticipating that these drugs would serve to competitively inhibit productive signaling interactions between sender and receiver cells. To test this possibility, we treated sender-receiver cell mixtures with biotin, biocytin, and bis-biotinamide (**Figure 2g**), measuring their effects on *trans*-cellular signaling following overnight incubation.

Among the three tested compounds, treatment with bis-biotinamide resulted in the most potent inhibition of trans-cellular signaling, with an approximate IC_50_ value of approximately 600 nM (**Figure 2h**). Treatment with biocytin also resulted in cell signaling inhibition, albeit with reduced efficacy relative to that of the multivalent bis-biotinamide. Free biotin was the least potent of the tested molecules, consistent with the binding preference of the immunoglobulin domain for derivatized species of the vitamin^40^. Taken together, these results indicate that *trans*-cellular interactions between our synthetic signaling components can be competitively blocked using soluble biotinamide derivatives. Notably, both biocytin and bis-biotinamide are cell-impermeant compounds^43^, and thus their effects are likely limited to extracellular components.

### Inducible signaling with bispecific receptor agonists

In addition to antagonizing AP-SynNotch, we also developed synthetic strategies in order to induce AP-SynNotch activation using exogenously administered agents. Toward this end, we devised two strategies: in a first approach, we created a bispecific “bridge” protein that could be used to induce *trans*-cellular complex formation between synthetic ligands and receptors^44^. Here, a monobiotinylated GFP (GFP-biotin) was generated by expressing a GFP-AP fusion protein within BirA-expressing cells. Following purification, the bispecific protein was added to mixtures containing anti-bio-ligand senders and receiver cells expressing an anti-GFP containing receptor. Whereas treatment with GFP-biotin induced productive signal activation, as determined via reporter detection, cultures treated with a non-biotinylated control GFP exhibited only background reporter levels, resembling that of non-treated controls (**Supplementary Figure 2c**).

In a second strategy, we tested a receptor in which the anti-biotinamide scFv served as the ECD, generating “anti-bio-SynNotch.” In this orientation, we reasoned that diverse biotin-containing compounds could be implemented in order to achieve divergent control of signaling activity, with individual molecules exhibiting either an antagonistic or agonistic property, depending upon its immobilization state (**Figure 3a**). To test the activity of the receptor, we evaluated anti-bio-SynNotch cells following overnight growth in microwells coated with biotinylated bovine serum albumin (biotin-BSA). Subsequent measurement of reporter expression levels revealed strong signaling activity in response to the immobilized biotinamide, confirming the inducibility of the receptor in this orientation (**Figure 3b**). Tests using a biotinidase-resistant^45^ biotin-BSA conjugate also resulted in anti-bio-SynNotch activation, as did desthiobiotin-BSA, albeit to a lesser extent as compared to biotinamide-based ligands (**Figure 3b, Supplementary Figure 3a-b**). In addition to immobilized biotins, coculture analyses using sender cells expressing an AP-containing ligand (AP-ligand) confirmed *trans*-cellular activity between anti-bio-SynNotch and AP-ligand, with the expression of BirA-KDEL being required of AP-ligand sender cells within this signaling configuration (**Figure 3c**).

**Figure 3.**
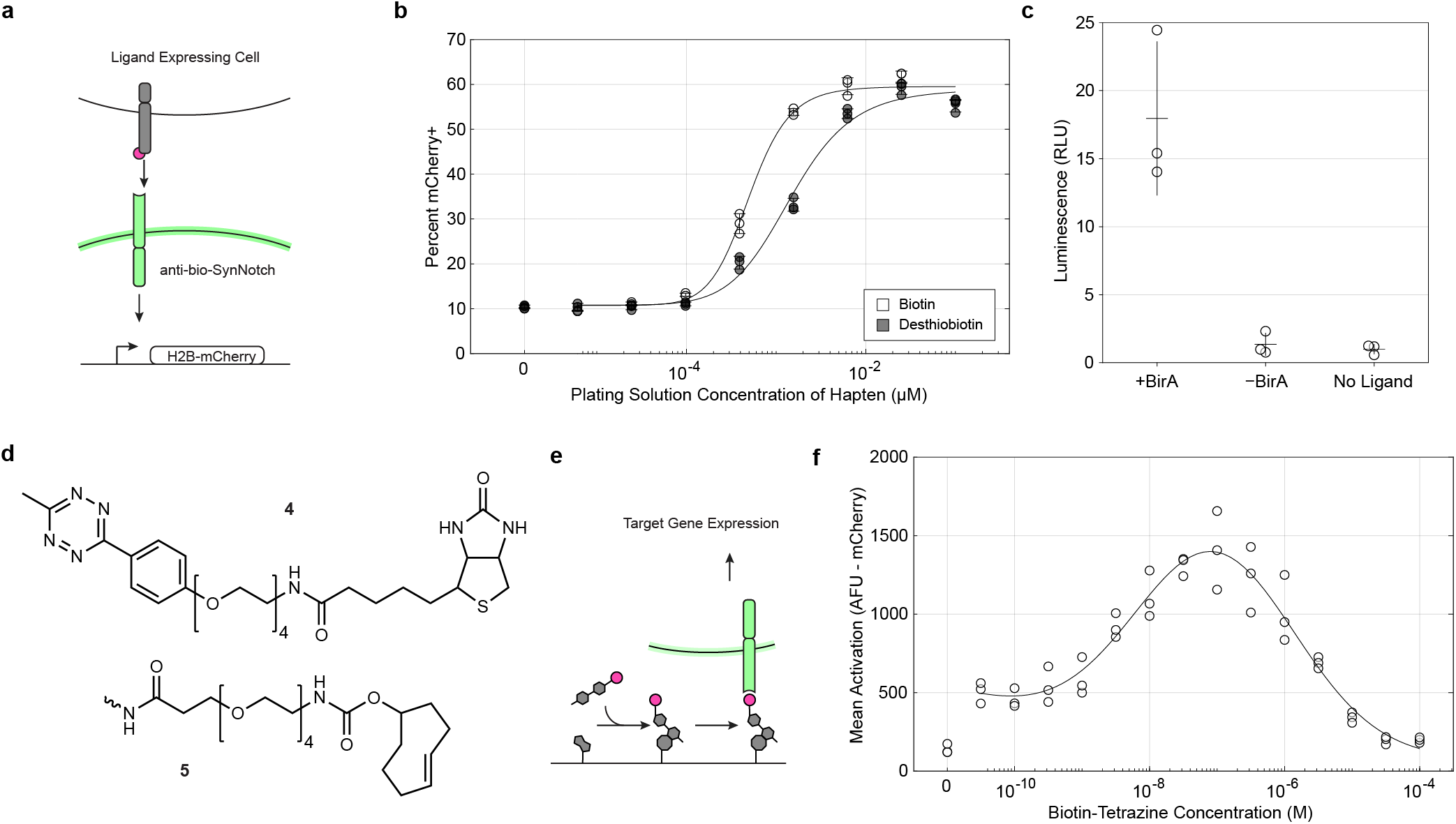
Signaling and chemical control of a biotinamide-binding receptor. **a**. Schematic depicting the trans-cellular activation of anti-bio-SynNotch by an AP-containing and BirA-modified ligand protein. **b**. Reporter levels from UAS-H2B-mCherry cells expressing anti-bio-SynNotch and stimulated with varying amounts of immobilized ligands based on biotinylated and desthiobiotinylated BSA conjugates. Error bars represent one standard deviation, n = 3 wells. **c**. Reporter levels from cocultures mixtures containing AP-ligand senders and anti-bio-SynNotch receiver cells. Receivers bearing a UAS-Luc reporter construct were tested against senders with and without coexpressed BirA-KDEL. Error bars represent one standard deviation, n = 3 wells. **d**. Structure of the conditional biotinamide-containing ligand biotin-tetrazine (4) and the reactive trans-cyclooctene (TCO) group (5), as conjugated to BSA. **e**. Schematic depicting the conversion of biotin-tetrazine into a signaling-competent (immobilized) ligand via bioorthogonal ligation with TCO-conjugated BSA. **f**. Reporter levels from UAS-H2B-mCherry cells expressing anti-bio-SynNotch and grown on surfaces containing immobilized TCO-BSA. Adherent cultures were treated with biotin-tetrazine at varying concentrations and mCherry levels were quantified 24 hours following biotin-tetrazine exposure. Sample size, n = 3 wells.

Using cells expressing anti-bio-SynNotch, we next devised a strategy in order to convert a soluble biotin-containing molecule into an immobilized (signaling competent) ligand. In this approach, we exploited a bioorthogonal ligation based on an inverse electron demand Diels-Alder reaction^46^ in order to immobilize a biotinamide-tetrazine compound to cell culture surfaces coated with fibronectin in combination with *trans*-cyclooctene (TCO) derivatized BSA (BSA-TCO, **Figure 3d-e**). Our anticipation was that the BSA-TCO could serve as an artificial extracellular matrix component, to which receptor-binding ligands could be attached. Indeed, cells that were grown in BSA-TCO coated microwells exhibited signaling responses that were dependent upon tetrazine-biotinamide treatment. Notably, tetrazine-biotinamide exhibited both agonistic and antagonistic activities, with a bell-shaped response to increasing doses of tetrazine-biotinamide (**Figure 3f**).

### PTM-dependent intracellular protein localization

Desiring to extend the utility of these tools to intracellular context, we next asked whether a synthetic, biotinamide binding domain could be used to recognize BirA-modified proteins within the cell interior. Here, we hypothesized conversion of the anti-biotinamide IgG to a single-chain Fab (scFab) format would permit its intracellular use, as has previously been demonstrated using other immunoglobulin sequences^47,48^ Thus, we generated an scFab by connecting the heavy and light chains of anti-biotinamide IgG via an 18 amino acid flexible linker^48^ resulting in “anti-bio-scFab.”

In order to test the functionality of the scFab, we carried out a fluorescence imaging-based analysis in which an mCherry-tagged immunoglobulin (anti-bio-scFab-mCherry) was coexpressed alongside a mitochondrially-targeted mTurquoise2 (mTurq2)-based BirA substrate protein (TOM20-mTurq2-3xAP) (**Figure 4a**). Here, we reasoned that the antigen-binding function of the scFab within the cytoplasm could be determined by measuring its extent of colocalization with biotinylated TOM20-mTurq2-3xAP. In transfected U-2 OS cells, fluorescence emissions from mCherry and mTurq2 were colocalized to the surface of mitochondria only in cells expressing cytoplasmic BirA (**Figure 4b-c**). In contrast, in cells lacking BirA, anti-bio-scFab-mCherry was diffusely distributed throughout the cytoplasm and nucleus. Thus, anti-bio-scFab is able to recognize biotinylated proteins within intracellular compartments, and our data provide evidence that BirA can be used to trigger PTM-dependent protein-protein complex formation within intracellular compartments.

**Figure 4.**
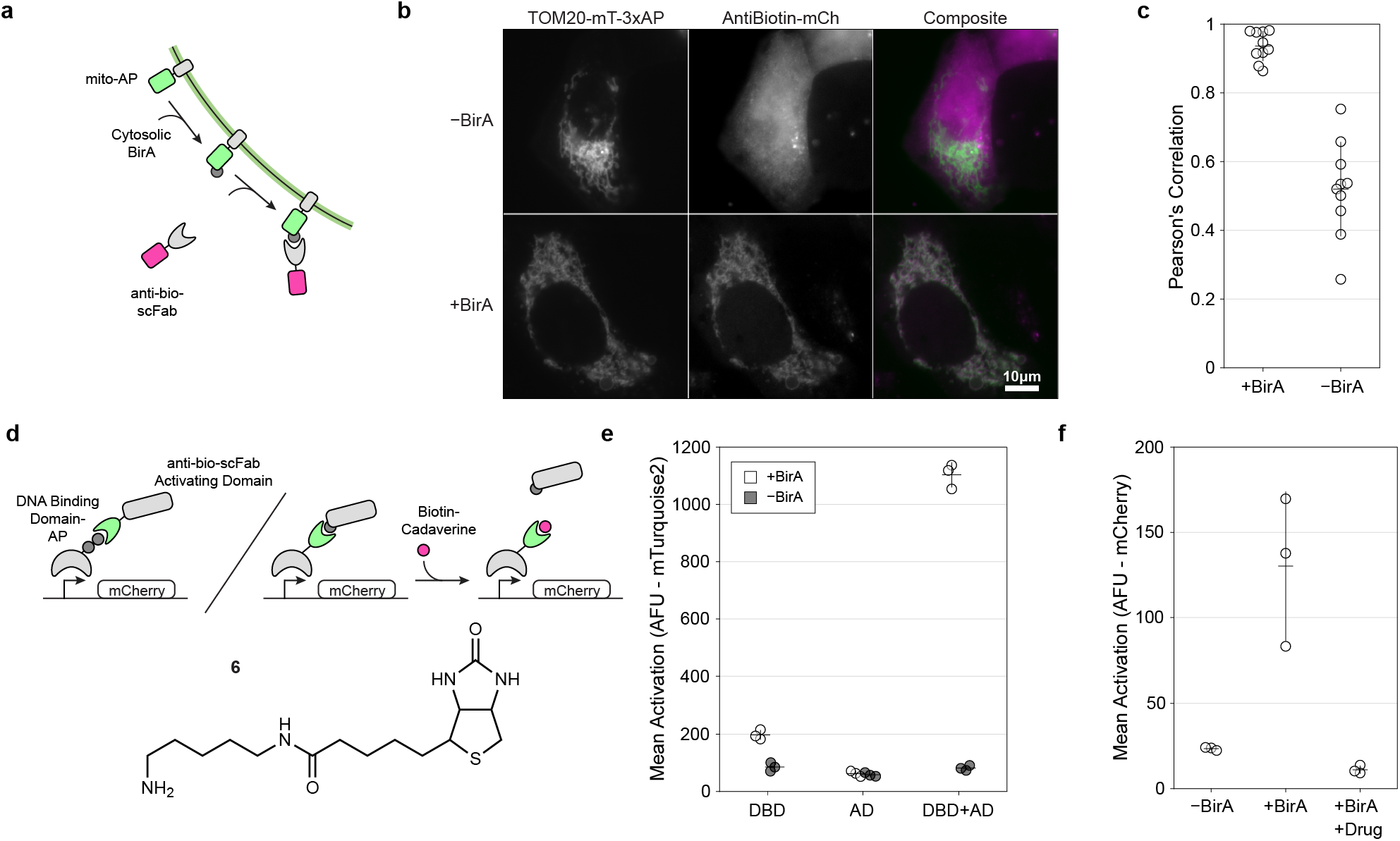
Design and validation of an scFab-based intracellular biotinamide-binding domain. **a**. Schematic depicting the colocalization assay used in the evaluation of an intracellular anti-bio-scFab. **b**. Representative fluorescence images of U-2 OS transfected with a bicistronic construct encoding TOM20-mTurq2-APx3 in combination with an IRES-driven anti-bio-scFab fused to mCherry. Localization between mTurq2 and mCherry fluorescence emissions were compared between cells that lacked (top) and contained (bottom) a coexpressed cytosolic BirA. **c**. Correlation coefficients of mTurq2 and mCherry fluorescence emissions in cells with and without cytosolic BirA. Values calculated using 10 representative images from each condition are displayed as Pearson’s coefficients. Error bars represent one standard deviation. **d**. Schematic depicting two methods of achieving PTM-dependent transcriptional activation. First, an AP-fused DNA Binding Domain which, when biotinylated, binds to anti-bio-scFab fused to an activating domain. Second, a Gal4 DBD fused to the anti-bio-scFab and its biotinylation induced interaction with an AP-VP64 activation polypeptide. The interaction can be disrupted with cell-permeant biotinamide containing molecules (top). The structure of the cell-permeant molecule biotin-cadavarine (6) (bottom). **e**. Gene activation levels from HEK293-derived TRE3G:mTurq2 reporter cells transfected expressing the TetR-APx2 DNA Binding Domain (DBD), anti-bio-scFab-p65-RTA activation domain domain (AD), or both, with and without coexpression of cytosolic BirA. Error bars represent one standard deviation, n = 3 wells. **f**. Reporter activation levels in HEK293 UAS:H2B-mCherry reporter cells expressing anti-bio-scFab-Gal4 and AP-VP64, with and without coexpression of cytosolic BirA, with comparison to BirA-expressing cells treated with 20 μM of biotin-cadaverine. Error bars represent one standard deviation, n = 3 wells.

### PTM-dependent transcriptional control

The functions of numerous intracellular proteins are controlled via PTMs, including that of Notch ICD (NICD) and its associated signaling proteins. For example, the nuclear translocation of the NICD requires ligand-induced sequential proteolysis of the receptor by ADAM10 (at a site within the NRR termed “S2”)^49^ and subsequently by the gamma-secretase complex (at a position within the TMD at “S3”)^50^. Additionally, the lifetime of the liberated ICD within the nucleus is determined via the phosphorylation of a C-terminally encoded degron^25,51^. Seeking to achieve PTM-mediated control over synthetic intracellular components, we next devised strategies to regulate nuclear proteins in a biotinylation-dependent manner.

Aiming to mimic the nuclear functions of the natural NICD, we sought to utilize BirA to gain PTM-dependent control over synthetic gene transcription. In this approach, we installed biotinylation-dependent control into the TRE3G promoter by generating a TetR DNA-binding domain fused to a tandem repeat of two AP fusion tags (TetR-2xAP). To quantify the activity to the transcription factor fusion, a TRE3G-regulated mTurq2 fluorescent reporter was added to the TetR-2xAP construct and used to measure levels of transcriptional output. In this design, we anticipated that modification of TetR-2xAP could be used to recruit transcriptional activation machinery to the TRE3G promoter, via a biotinylation-induced interaction with anti-bio-scFab fused to p65 and RTA domains (scFab-p65-RTA, **Figure 4d**). To test this hypothesis, we coexpressed the TetR-2xAP-TRE3G-mTurq2 construct and scFab-p65-RTA in HEK293 cells with and without cytosolic BirA. Indeed, cells in which all three components were co-expressed led to the greatest mTurq2 expression, whereas cells lacking BirA had expression levels comparable to cells lacking the reporter (**Figure 4e**). In a similar design, a fusion between the anti-bio-scFab and the Gal4 DNA binding domain (anti-bio-scFab-Gal4) was used in combination with an AP fused version of the transcriptional activator VP64 (AP-VP64). In transfected UAS:H2B-mCherry reporter cells, these components were also able to facilitate biotinylation-dependent reporter expression (**Figure 4f**). Together, these data demonstrate that BirA biotinylation can be utilized to regulate the formation of transcriptional complexes for gene expression control.

Lastly, we asked whether intracellular transcriptional complexes could be disrupted using exogenously applied biotinamide. To test this possibility, we treated BirA-positive cells with biotin-cadaverine, a membrane-permeant analog of biocytin^52^. Measurement of reporter levels in BirA-positive UAS:H2B-mCherry cells showed diminished amounts of H2B-mCherry expression following biotin-cadaverine treatment, indicating that the formation of active transcription complexes can be competitively blocked in a dose-dependent manner using biotin-cadaverine (**Figure 4f, Supplemental Figure 4**). Thus, control of PTM-dependent signaling systems can be achieved via the membrane-permeant biotin-cadaverine.

## DISCUSSION

In summary, we have described a chemogenetic toolkit for installing PTM-based control into synthetic biological systems. Drawing inspiration from natural regulatory mechanisms, we constructed an encodable “writer/reader” framework, leveraging *E. coli* BirA as an orthogonal “writer” module for modifying synthetic signaling proteins containing the AP substrate tag. To enable the programming of cellular activities in response to PTM events, we designed synthetic “reader” elements via a biotinamide specific antibody and used fusion proteins containing the domain in order to facilitate PTM-dependent functions.

In addition, we demonstrated the ability to modulate cell-cell communication by exerting genetic regulation over the expression of writer and reader elements. Through control over the expression of the receptor, as well as its modifying enzyme, we were able to modulate the receptor’s specificity and signaling receptivity to various ligands. In addition, through the use of externally-administered drug molecules, we exerted chemical control over the assembly state and signaling activity of the encoded systems via both naturally-occurring as well as synthetic biotinamide-containing compounds. Furthermore, we implemented the biotinylation strategy as a reader/writer framework for achieving PTM-dependent regulation within intracellular contexts, validating the biotinamine binding capability of a cytoplasmically-localized scFab and demonstrating its utility in achieving synthetic gene expression control.

The present work validates the utility of BirA-mediated PTM in the design of synthetic regulatory systems in mammalian cells. Biotinylation is a rare PTM with respect to mammalian proteomes, with only one known human cytoplasmic protein bearing the modification (acetyl-coA carboxylase), four that reside within mitochondria, and none far which have been identified as localized to the plasma-membrane^53^. Additional engineering at the level of the protein ligase will likely elaborate the synthetic potential of the platform future applications. For example, chemogenetic and optogenetic control over BirA may provide more precise ways to regulate the timing and spatial localization of PTM events, and further evolution of BirA may enable the ligation of new PTMs^54–57^. An advantageous feature of the current system – and one that may provide unique synthetic utility – is the ability of the reader domain to recognize biotinamide, a safe and naturally occurring compound. In our studies, we exploited this recognition in order to achieve control over the PTM-mediated protein-protein interactions using readily-available molecules, including naturally occurring vitamins and cofactors (biotin and biotinamide), as well as their synthetic derivatives (bis-biotinamide, tetrazine-biotin).

Finally, we note that a motivating rationale for the development of these tools is to enable the elaboration of synthetic signaling pathways in ways that more closely mimic the intricacies of natural systems. Indeed, it is anticipated that the sophistication of synthetic capabilities – especially regarding the development of complex multicellular systems – will emerge. We hope that these systems, or others, will contribute to these advances. Throughout natural systems, PTMs play a prominent and critical role, exhibiting the benefits of rapid tunability as a secondary control system. Therefore, we anticipate that the incorporation of deliverable and tunable PTM control strategies will be a necessary and central component in the realization of future synthetic systems.

## METHODS

### Plasmid Construction

Standard cloning procedures were used in the generation of all DNA constructs. DNA fragments were amplified with Phusion High-Fidelity DNA Polymerase (New England Biolabs), and Gibson assembly was accomplished using the NEBuilder HiFi DNA Assembly master mix (New England Biolabs). New England Biolabs restriction enzymes were used to digest DNA, and T4 DNA Ligase (New England Biolabs) was used for ligation. The pDisplay-BirA-ER construct (Addgene #20856) was a gift from Alice Ting (Stanford University). The pLV-EBFP2-nuc construct (Addgene #36085) was a gift from Pantelis Tsoulfas. The GAL4UAS-Luciferase reporter was a gift from Moritoshi Sato (Addgene #64125). *Sequence files and the plasmids described in this work have been deposited to AddGene and are available under via following depositions: X - #####, Y - #####, Z - #####*.

### Mammalian Cell Culture

Mammalian cell lines were cultured in a humidified incubator maintained at 37 °C with 5% CO_2_. HEK293FT cells (ThermoFisher) and U2OS (Sigma-Aldrich) cell lines were cultured in high-glucose DMEM containing sodium pyruvate, and supplemented with 10% FBS, nonessential amino acids (LifeTechnologies), Glutamax (LifeTechnologies), Penicillin-Streptomycin (50 units-μg/mL; Gibco). HEK293FT cells were maintained in G418-containing media (500 μg/mL; Invivogen). Experiments involving doxycycline-inducible, or doxycycline-sensitive transcription factors (rtTA-3 and TetR-NS3-VP64, respectively) were carried out using a growth medium containing 10% Tet-Approved FBS (Clontech) in place of standard growth serum. Stable cell lines generated using DNAs encoding antibiotic resistance markers, and stably integrated cells were selected and maintained in media containing either zeocin (100 μg/mL; Invivogen), puromycin (500 ng/mL; Invivogen), Hygromycin B-Gold (75 μg/mL; Invivogen), blasticidin (10 μg/mL; Invivogen), or combinations thereof, as appropriate per cell line (see Supplementary Table 1).

### DNA Transfection

DNA transfections were carried out using Lipofectamine 3000 Reagent (ThermoFisher) according to the manufacturer’s protocol. For coculture assays, luciferase reporter plasmids were reverse transfected by first adding the transfection mixture to each well for at least 15 min, and then adding the receptor-expressing cells in media.

### Stable Cell Line Generation

HEK293FT cells were grown in 6-well plates and cotransfected with plasmid mixtures containing a lentiviral packaging vector (psPAX2), an envelope-encoding plasmids vectors (VSV-G), and a transfer plasmid encoding relevant transgenes. Viral particles were harvested at 24 hr and 48 hr times points after transfection, and cellular debris were removed from viral supernatants via centrifugation (300 x G, 5 min) followed by filtration through 0.45 μm porous membrane. Viral containing media was either used for transduction immediately or stored at −80 °C for later use. For transduction, virus-containing media was diluted into culture wells containing the indicated cell lines. Viral transductions were allowed to proceed for 48 hr under growth conditions prior analysis, or selection using the antibiotic concentrations indicated above. The appropriate antibiotic was added for ten days, and single clones were isolated via limited dilution. For cell lines established using pcDNA3.1 derived vectors, cells were transfected with linearized plasmid and selected with antibiotic at 48 hr post-transfection. Clone isolation via limited dilution in 96-well plates was performed as described above.

### Western Blots

Cell lysates were prepared by direct lysis in RIPA Lysis and Extraction Buffer (Alfa Aesar) followed by sonication. Proteins were separated by denaturing polyacrylamide gel electrophoresis using NuPAGE SDS-PAGE gels (Thermo Fisher). Proteins were transferred to nitrocellulose membranes and blocked with 5% nonfat dry milk prior to probing with streptavidin (using a direct conjugate with HRP), or a mouse anti-myc antibody (followed by detection with anti-mouse HRP). Chemiluminescent detection of HRP was carried out using the SuperSignal West Pico PLUS Chemiluminescent Substrate (Pierce).

### Fluorescence Microscopy

Cells were imaged by epifluorescence microscopy after having been plated on 8-well Optically Clear Plastic Bottom slides (Ibidi) coated with Fibronectin. During imaging, cells were maintained in PBS or standard culture media. For immunofluorescent staining of fixed cells, cells were fixed for 10min at room temperature with paraformaldehyde (4% v/v in PBS from 16% solutions purchased from Thermo Fisher) and rinsed with PBS. Cells were blocked with a BSA solution (5% w/v in PBS) before being incubated with fluorophore conjugated streptavidin or antibody. Images were acquired with ZEN imaging software (Zeiss). Image files were processed with a custom MATLAB (Mathworks) script in order to adjust contrast uniformly across experimental conditions. Pearson’s coefficients were determined by measuring pixel intensities within an mCitrine positive mask for 10 images in each condition with a MATLAB (Mathworks) analysis script (custom script; available upon request).

### Plated Ligand Assay

Non tissue-culture treated 96-well plates were coated with immobilized ligands using a coating solution containing fibronectin (5 ng/mL) and ligand protein at an appropriate concentration. Solutions prepared in PBS were dispensed into individual wells using 50 μL volumes; coating reactions were then allowed to proceed for 1 hr at room temperature. Wells were rinsed three times with 200 μL PBS, and receptor-expressing receiver cells were then seeded within individual wells in growth medium at a density of 4 × 10^4^ cells per well. Receptor activation was measured 24 hr post-plating using either epifluorescence microscopy or flow cytometry (for cells containing fluorescence-based reporter constructs; see ‘Flow Cytometry’ section below), or chemiluminescent detection using (for cells containing firefly luciferase (fLuc)-based reporter constructs, see ‘Coculture Luciferase Assay’ section below), as indicated within the text and figure captions. Anti-biotin IgG (clone Bio3-18E7) was from Miltenyi Biotech. Mouse anti-myc IgG was from (ThermoFisher). Initial experiments using serial dilutions of ligands were used to determine appropriate coating solution working concentrations; 1 ng/mL to 1 μg/mL for antibody-based ligands, and 10 pM to 100 nM for ligands based on BSA-conjugates.

### Flow Cytometry

Flow cytometry analyses were carried out using an AttuneNxT instrument (ThermoFisher), and detection parameters were set with gating for single cells by scatter detection (**Supplementary Figure 5a-c**). Output files were analyzed using a MATLAB (Mathworks) analysis script (custom script; available upon request). The percent activation was determined by calculating the percentage of cells with mCherry expression levels above a determined threshold using the non-transfected control as a guide.

### Coculture Luciferase Assay

The luciferase assay used for coculture studies was adapted from Gordon *et al*^4^. Synthetic Notch receiver cells were reverse transfected in a 96-well plate (5 × 10^4^ cells per well) with 9.9 ng of UAS:Firefly-Luciferase plasmid and 0.1 ng Nanoluc plasmid per well. At 24 hr post-transfection, the media for each well was replaced with fresh media containing 8 × 10^4^ sender cells. 48 hr post-transfection, cells were lysed (Nano-Glo Dual-Luciferase Reporter Assay System, Promega) and the luminescence was measured according to the manufacturer’s protocol. fLuc luminescence activity levels were normalized to that of nanoluciferase, as introduced to cells using a constitutively expressed nanoluciferase transfection control plasmid, provided by the manufacturer, and measured according to the manufacturer’s protocol.

### Protein Conjugation

Bovine serum albumin (BSA) was conjugated to biotin, or desthiobiotin using activated *N*-hydroxysuccinimidyl (NHS) esters. Reactions were carried out in solutions containing 75 μM BSA dissolved in 75 mM sodium bicarbonate buffer (pH 8.2). Protein conjugation reactions were initiated by the addition of NHS esters from stock solutions dissolved in DMSO. Final reaction solutions contained 7.5 mM of each NHS ester and 20% (v/v) DMSO. Coupling reactions were allowed to proceed for 1 hour at room temperature, followed by removal of unconjugated esters via three rounds of dialysis against PBS using 3000, or 10000 MWCO dialysis membranes. Following dialysis, the solution was sterilized via filtration through a 0.2 μm porous membrane. Extents of conjugation for TAMRA-containing conjugates was determined via the extinction coefficient (92000 M^-1^cm^-1^ at 555 nm for TAMRA). For BSA conjugates not linked to chromophores, extent of biotin-conjugation was determined using a 4’-hydroxyazobenzene-2-carboxylic acid (HABA)-based assay (Biotin Quantitation Kit, Pierce) according to the manufacturer’s protocol. A BSA conjugate containing a biotinidase-resistant biotin derivative (biotinoyl-2-aminobutyric acid), as well as BSA containing *trans*-cyclooctene functionality, were generated according to the same procedure using NHS-dPEG4-biotinidase resistant biotin (Quanta Biodesign) and TCO-PEG4-NHS ester (Click Chemistry Tools), respectively.

### Protein Expression and Purification

DNA sequences encoding a GFP-AP fusion, or a control GFP containing a mutated AP sequence (GLNDIFEAAAAEWHE), were cloned into the pBAD bacterial expression vector with a N-terminally encoded 6xHis-tag. Expression was carried out in BirA-containing *E. coli* cells (AVB101, Avidity) grown in lysogeny broth. Cells were grown until culture density OD_600_ values reached 0.4 −0.6, at which point expression was induced by the addition of L-arabinose to a final concentration of 0.2% (w/v), IPTG to a concentration of 1.5 mM, and d-biotin to a concentration of 50 μM. Following expression for 3 h at 30 °C, cells were harvested via centrifugation and stored as frozen pellets at −20 °C until purification. Metal-chelate affinity chromatography was used to purify the 6xHis-tagged proteins using the Ni-NTA Fast Start Kit (Qiagen), according to the manufacturer’s protocol. Following purification, imidazole was resolved from protein solutions by dialysis against PBS using procedures similar to that described above for BSA.

### Compounds

Stock solutions of biotin and biocytin were stored in water at 10 mM at −20 °C. Stock solutions of biotin-tetrazine (biotin-PEG4-methyltetrazine, Click Chemistry Tools), bis-biotinamide (bis-dPEG3-biotin, Quanta Biodesign) and biotin-cadaverine (ThermoFisher) were stored in DMSO at 10 mM at −20 °C. Grazoprevir was stored in DMSO at 10 mM at −20 °C and used at a working concentration of 5 μM, added at the time of transfection.

## ACKNOWLEDGEMENTS

This work was supported through funding from the NIH NIGMS (GM128859). JBM was supported through a Kilachand Graduate Fellowship awarded by the Multicellular Design Program at Boston University.

## AUTHOR CONTRIBUTIONS

All authors designed and carried out experiments, analyzed data, and contributed to the preparation of the manuscript.

## COMPETING INTEREST STATEMENT

None

## Notes

### Competing Interest Statement

JTN is an inventor on a patent related to the work described herein.

