## Supplementary Information for "A Genetically Encodable and Chemically Disruptable System for Synthetic Post-Translational Modification Dependent Signaling"

**Supplementary Figure 1: Biotinylation and signaling activity of AP-SynNotch.**

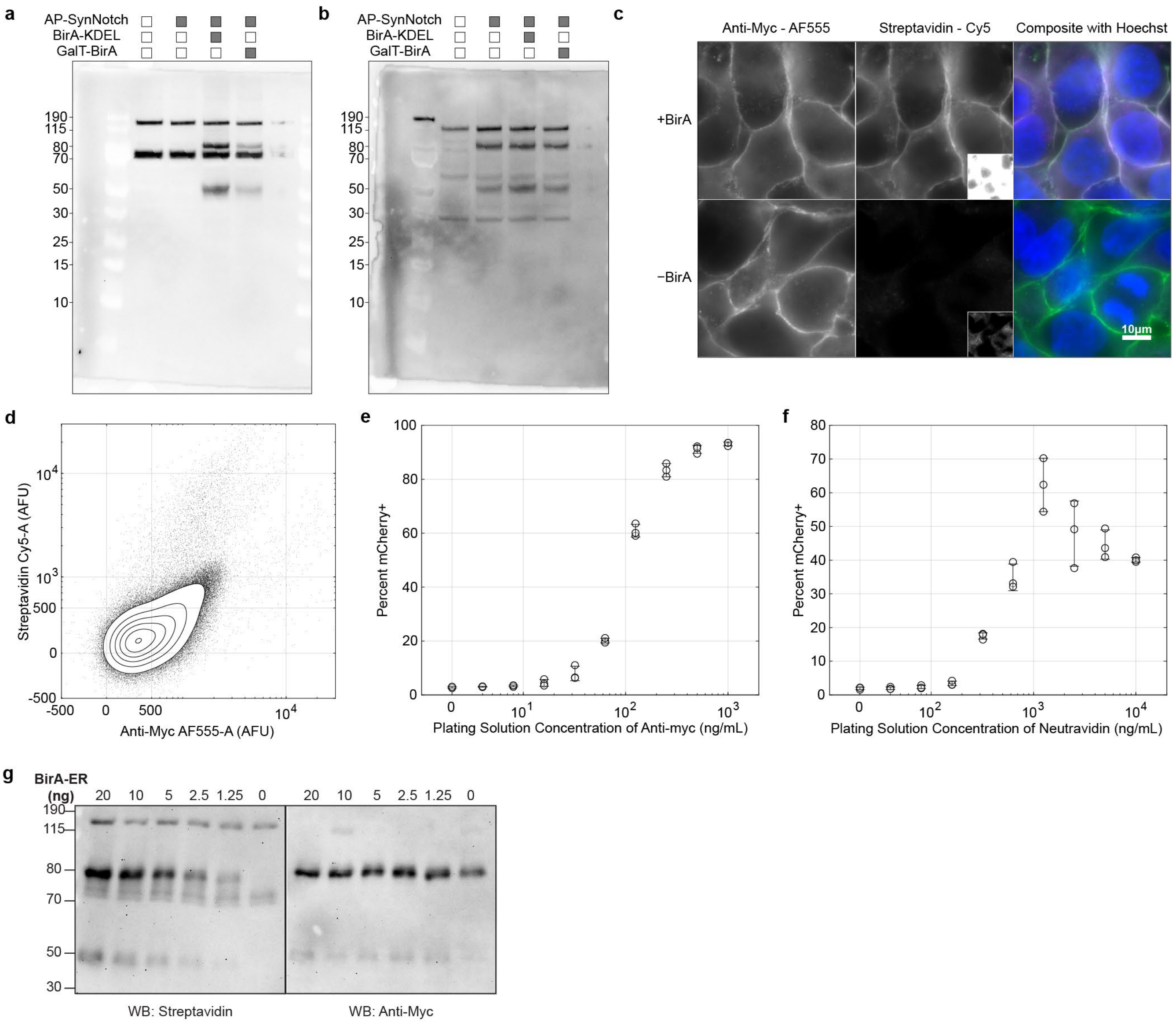


**a.** Uncropped and unadjusted Western blot from Figure 1b for HEK293 cells expressing AP-SynNotch (Well 2), and co-expressing BirA-KDEL (Well 3) or GalT-BirA (Well 4). Non-transfected is Well 1. The Western Blot was probed with Streptavidin-HRP. **b.** Uncropped and unadjusted Western blot from Figure 1b HEK293 cells expressing AP-SynNotch (Well 2), and co-expressing BirA-KDEL (Well 3) or GalT-BirA (Well 4). Non-transfected is Well 1. The Western Blot was probed with anti-myc-HRP. **c.** AP-SynNotch is biotinylated by BirA-KDEL and presented at the cell surface. **d.** Non-transduced HEK293 cells were probed with streptavidin-Cy5 and anti-myc-AF555 and the fluorescence was quantified via flow cytometry. The sample includes 120,401 cells. **e.** AP-SynNotch expressing cells were cultured on wells coated with different concentrations of anti-myc antibody. Error bars represent one standard deviation, n = 3 wells. **f.** AP-SynNotch expressing cells co-expressing BirA-KDEL were cultured on wells coated with different concentrations of Neutravidin. Error bars represent one standard deviation, n = 3 wells. **g.** HEK293 cells were transfected with a construct containing AP-SynNotch at 10 ng per well and co-transfected with different amounts of a construct containing BirA-KDEL.

**Supplementary Figure 2. Cell-mediated trans-activation and chemical regulation of signaling from biotinylated AP-SynNotch receptors.**

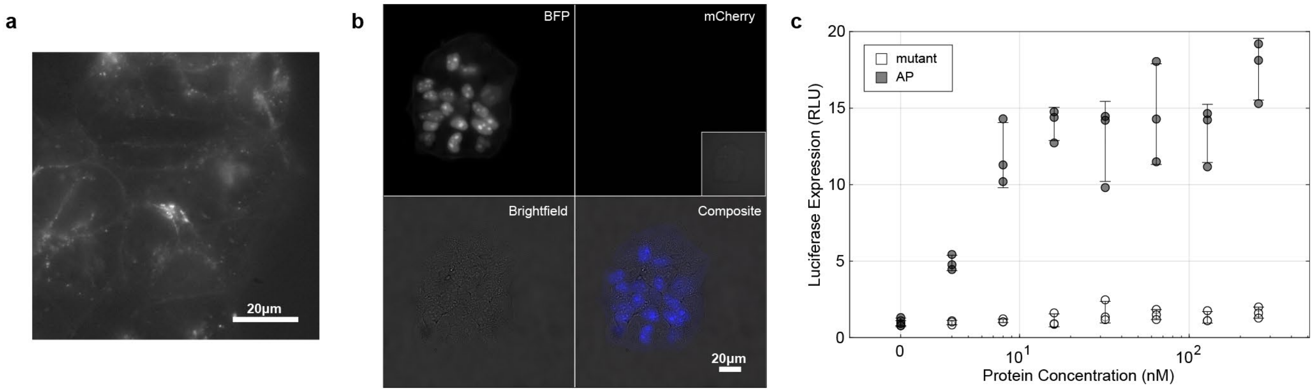


**a.** Labeling of anti-bio-ligand expressing HEK293 cells with SNAP-Surface-AF647 indicates membrane localization of the ligand. Internalized punctae signify the ability for the ligand to generate endocytic force. **b.** AP-SynNotch expressing cells that co-express BirA-KDEL and nuclear BFP do not trigger activation of downstream mCherry expression when they are not interacting with anti-bio-ligand expressing cells. Inset indicates increased contrast. The composite image indicates BFP (blue), mCherry (red), and Brightfield (white). **c.** AP-SynNotch cells, which contain an extracellular Anti-GFP nanobody, were cocultured with anti-bio-ligand expressing cells. Purified GFP, either containing AP or a mutated AP sequence in which the modified lysine and its adjacent residues were mutated to alanine (GLNDIFEAAAAEWHE), which was expressed in a BirA expressing *E. coli* strain, was added to wells at different concentrations. Error bars represent one standard deviation, n = 3 wells.

**Supplementary Figure 3. Signaling and chemical control of a biotinamide-binding receptor.**

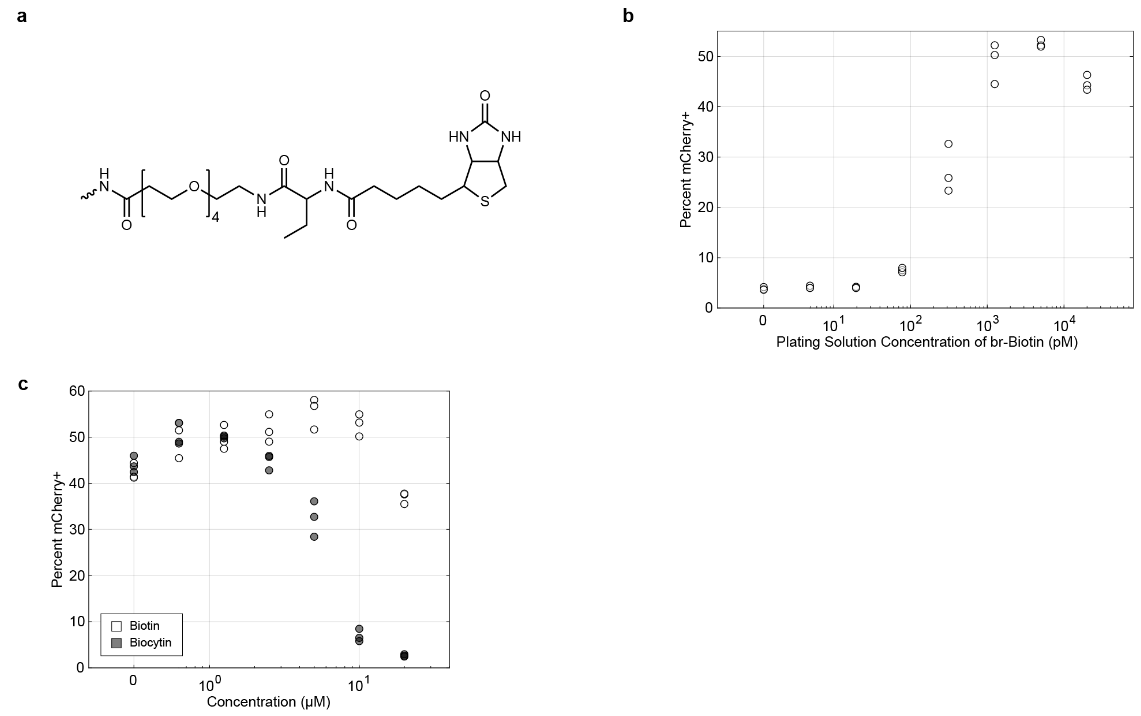


**a.** Biotinidase-resistant biotin-NHS ester was conjugated to BSA. The ethyl group adjacent to the amide group of biotinamide prevents the hydrolysis and release of biotin by Biotinidase. **b.** Anti-bio-SynNotch expressing cells are activated when cultured on wells coated with Biotindase-resistant (br) biotin-conjugated BSA. Number of samples: n = 3 wells. **c.** Anti-bio-SynNotch expressing cells cultured on biotin-BSA were cultured at time of plating with different concentrations of biotin and biocytin. Biocytin is more effective at competitively inhibiting the antibody fragment. Number of samples: n = 3 wells.

**Supplementary Figure 4. Design and validation of an scFab-based intracellular biotinamide-binding domain.**

a

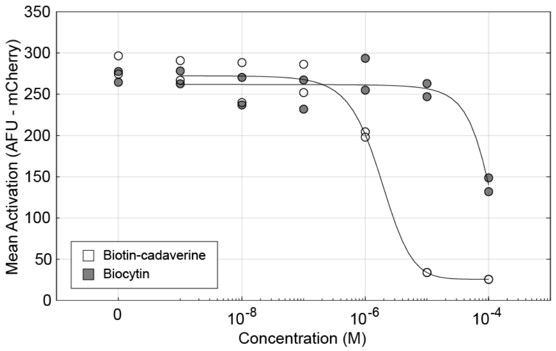


**a.** Reporter HEK293 cells with an integrated UAS:H2B-mCherry construct were transfected with a construct containing constitutive expression of AP-VP64 and anti-bio-scFab-Gal4. The cells were cotransfected with a construct containing Citrine-HA-BirA. At the time of transfection, samples were incubated with different concentrations of biocytin and biotin-cadaverine. Samples were measured the next day with flow cytometry, gating for Citrine expression. Number of samples: n = 2 wells.

**Supplementary Figure 5: Gating Schemes for Flow Cytometry**

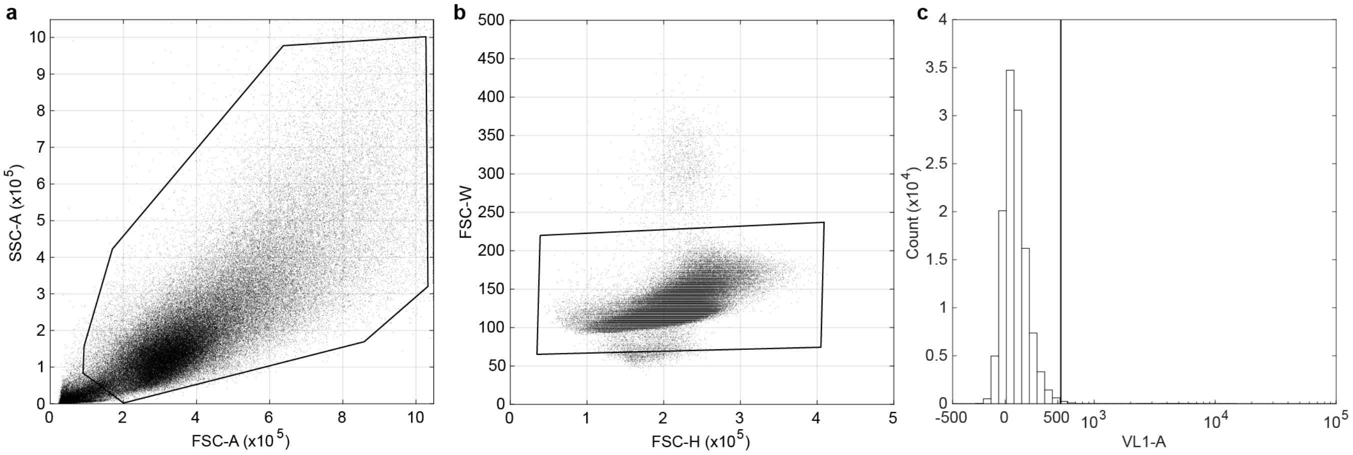


**a.** Cells are first gated in a Forward vs. Side Scatter plot to eliminate dead cells and debris. **b.** Cells are then gated for singlets through a Forward Scatter Height vs. Width gate. **c.** Finally, if cells have a transfection marker, they are gated positive for that fluorescent marker. The gate is adjusted to only include about 1% of cells which do not express the fluorescent marker.

| Cell Line | Source/Derived From | Selection Antibiotic |
|---|---|---|
| HEK293FT | ThermoFisher (catalog #R70007) | Geneticin |
| UAS Reporter (UAS:H2B-mCherry) | HEK293FT | Geneticin, Zeocin |
| AP-SynNotch | UAS Reporter (UAS:H2B-mCherry) | Geneticin, Zeocin, Hygromycin |
| AP-SynNotch + BirA-ER | AP-SynNotch | Geneticin, Zeocin, Hygromycin, Blasticidin |
| AP-SynNotch + T3G:BirA-ER | AP-SynNotch | Geneticin, Zeocin, Hygromycin, Puromycin |
| anti-bio SynNotch | UAS Reporter (UAS:H2B-mCherry) | Geneticin, Zeocin, Hygromycin |
| anti-bio-ligand | HEK293FT | Geneticin, Puromycin |
| U-2 OS | Sigma-Aldrich (catalog #92022711) | n/a |

**Supplementary Table 1. Sources of the cell lines used and generated in this study.** Antibiotic concentrations used in the selection and maintenance of the indicated cell lines are described in the Methods.
